# Pseudogenes in the mouse lineage: transcriptional activity and strain-specific history

**DOI:** 10.1101/386656

**Authors:** Cristina Sisu, Paul Muir, Adam Frankish, Ian Fiddes, Mark Diekhans, David Thybert, Duncan T. Odom, Paul Flicek, Thomas Keane, Tim Hubbard, Jennifer Harrow, Mark Gerstein

## Abstract

Pseudogenes are ideal markers of genome remodeling. In turn, the mouse is an ideal platform for studying them, particularly with the availability of developmental transcriptional data and the sequencing of 18 strains. Here, we present a comprehensive genome-wide annotation of the pseudogenes in the mouse reference genome and associated strains. We compiled this by combining manual curation of over 10,000 pseudogenes with results from automatic annotation pipelines. Also, by comparing the human and mouse, we annotated 165 unitary pseudogenes in mouse, and 303 unitaries in human. We make all our annotation available through mouse.pseudogene.org. The overall mouse pseudogene repertoire (in the reference and strains) is similar to human in terms of overall size, biotype distribution (~80% processed/~20% duplicated) and top family composition (with many GAPDH and ribosomal pseudogenes). However, notable differences arise in the pseudogene age distribution, with multiple retro-transpositional bursts in mouse evolutionary history and only one in human. Furthermore, in each strain about a fifth of the pseudogenes are unique, reflecting strain-specific functions and evolution. Additionally, we find that ~15% of the pseudogenes are transcribed, a fraction similar to that for human, and that pseudogene transcription exhibits greater tissue and strain specificity compared to protein-coding genes. Finally, we show that highly transcribed parent genes tend to give rise to processed pseudogenes.

## Introduction

The mouse is one of the most widely studied model organisms [1], with the field of mouse genetics accounting for more than a century of studies towards understanding mammalian physiology and development [2, 3]. Recent advances of the Mouse Genome Project [4, 5] towards completing the *de novo* assembly and gene annotation of a variety of mouse strains and the plethora of developmental and transcriptional data available from the mouse ENCODE project, provide a unique opportunity to get an in-depth picture of the evolution and variation of these closely related mammalian organisms.

Mice frequently have been used as a model organism for studying human diseases due to their experimental tractability and similarities in their genetic makeup with humans [6]. Scientists have achieved this by developing mouse models of specific diseases and generating knockout mice to recapitulate phenotypes associated with a loss-of-function (LOF) mutation observed in humans. The advent of high-throughput sequencing has led to the emergence of population and comparative genomics as new windows into the relationship between genotype and phenotype among the human population. Current efforts to catalog genetic variation among closely related mouse strains extend this paradigm.

Since their divergence around 90 million years ago (MYA) [7, 8, 9, 10, 11] the human and mouse lineages followed a parallel evolutionary pattern [12]. Although it is hard to directly compare the two species, there is a large range of divergence in the mouse lineage, with some approaching human-chimp divergence levels in terms of the number of intervening generations [12] (**Figure 1A**). The mouse strains under investigation have differences in their genetic makeup that manifest in an array of phenotypes, ranging from coat/eye colour to predisposition for various diseases [5]. Moreover, the generation of these strains has been extensively documented [13]. Following a well-characterised inbreeding process for at least 20 sequential generations, the inbred mice are homozygous at nearly all loci and show a high level of consistency at genomic and phenotypic levels [14]. This helps minimise a number of problems raised by the genetic variation between research animals [15]. repeated inbreeding has also resulted in substantial differences between mouse strains, giving each strain the potential to react uniquely to an acquired mutation [16].

**Figure 1.**
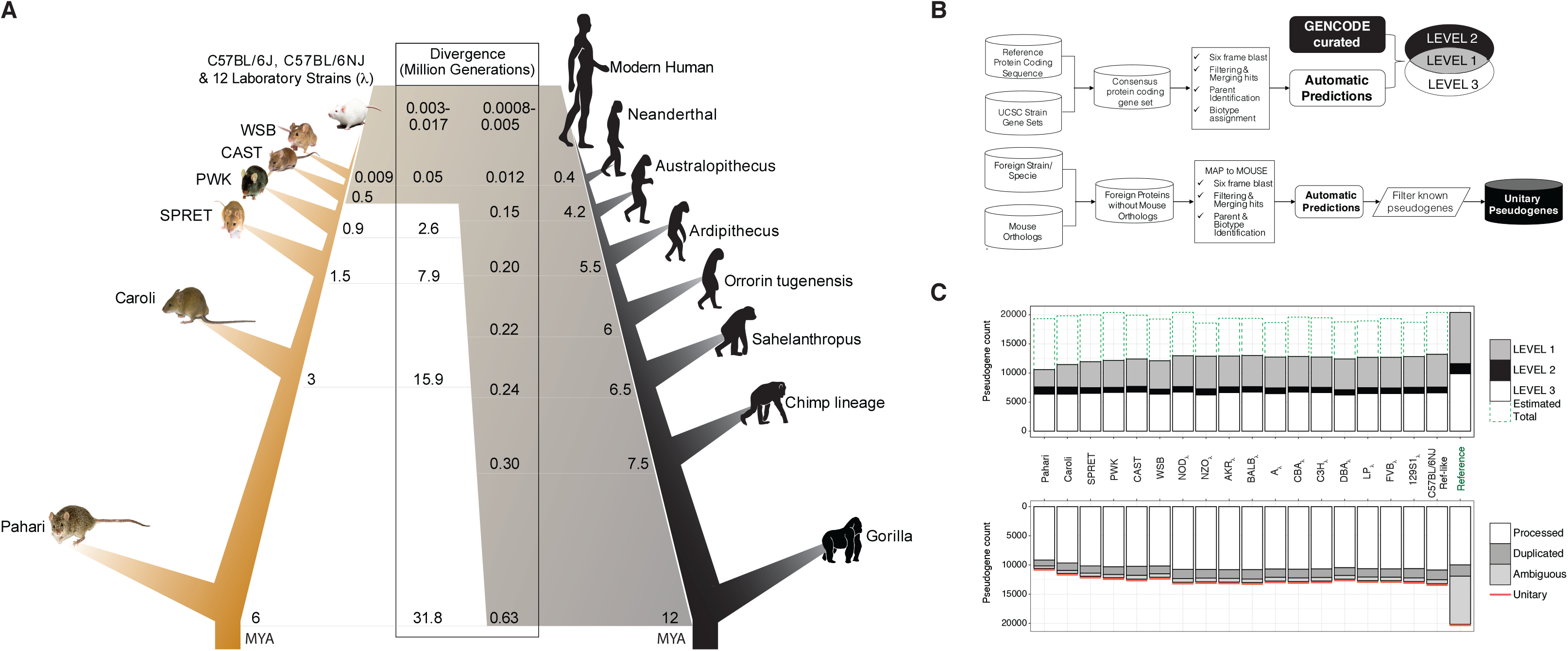
A – Human vs. mouse lineage comparison. *MYA* – million years ago, l – laboratory strain. B (top) – pseudogene annotation workflow for mouse strains. B (bottom) – unitary pseudogene annotation pipeline. C – Summary of mouse strains’ pseudogene annotation. *Level 1* are pseudogenes identified by automatic pipelines and liftover of manual annotation from the reference genome; *Level 2* are pseudogenes identified only through the liftover of manually annotated cases from the reference genome; *Level 3* are pseudogenes identified only by the automatic annotation pipeline.

To uncover key genome remodeling processes that govern mouse strain evolution, we focused on the pseudogene complements of each strain, while highlighting their shared features with the human genome. In this paper, we describe the first pseudogene annotation and analysis of 18 widely used inbred mouse strains alongside the reference mouse genome. Additionally, we provide the latest updates on the pseudogene annotation for both the mouse and human reference genomes, with a particular emphasis on the identification of unitary pseudogenes with respect to each organism.

Often regarded as genomic relics, pseudogenes provide an excellent perspective on genome evolution [17, 18, 19, 20]. Pseudogenes are DNA sequences that contain disabling mutations rendering them unable to produce a fully functional protein. Different classes of pseudogenes are distinguished based on their creation mechanism: (i) processed pseudogenes, formed through retrotransposition, (ii) duplicated pseudogenes, formed through gene duplication and subsequent disablement of one of the duplicates, and (iii) unitary pseudogenes, formed when functional genes acquire disabling mutations that result in the inactivation of the original coding loci. Unitary pseudogenes are also characterised by the presence of a functional orthologous gene on the same locus in other species. Additionally, pseudogenes that are present in a population as both functional and nonfunctional alleles are termed polymorphic [21]. Such pseudogenes represent disablements that have occurred on a much more recent timescale. These are LOF mutations that are not fixed in the population and are still subject to evolutionary pressures [21]. From a functional perspective, pseudogenes can be classified into three categories: (i) *dead-on-arrival*, which are elements that are nonfunctional and are expected to be eliminated from the genome with time, (ii) *partially active*, which exhibit residual biochemical activity, and (iii) *exapted*, which are elements that have acquired new functions and can interfere with the regulation and activity of protein-coding genes.

Moreover, pseudogenes reflect changes in selective pressures and genome remodeling forces. Duplicated pseudogenes can reveal the history of gene duplication, one of the key mechanisms for establishing new gene functions [22]. While the majority of duplicated gene copies are eventually pseudogenised [23], successfully retained paralogs can acquire new functions [24], a process known as neofunctionalisation [25]. Furthermore, duplicated pseudogenes can help explore the role of gene dosage in the inactivation or preservation of duplicated genes [26, 27]. Processed pseudogenes inform on the evolution of gene expression as well as the history of transposable element activity, whereas unitary pseudogenes are indicative of gene families that died out. Thus, pseudogenes can play an important role in evolutionary analysis as they can be regarded as markers of LOF events.

An LOF event is a mutation that results in a modified gene product that lacks the molecular function of the ancestral gene [28]. Unitary pseudogenes are an extreme case of LOF, where mutations that result in complete inactivation of a gene are fixed in the population. In recent years, LOF mutations have become a key research topic in genomics. In general, loss of a functional gene is detrimental to an organism's fitness. A number of studies have showcased the evolutionary advantages of the accumulation and fixation of LOF mutations that result in the formation of unitary pseudogenes [29, 30, 31, 32]. One notable example is the pseudogenisation of proprotein convertase subtilisin/kexin type 9 (PCSK9) in human evolution. This is commonly associated with reduced risk of heart disease by lowering plasma low-density lipoprotein (LDL) levels. This is achieved by preventing the production of PCSK9 protein and its subsequent binding to and degradation of cellular LDL receptors [33]. By contrast, gain-of-function (GOF) mutations that revert the disablement and result in the production of PCSK9 are commonly associated with an enrichment in plasma LDL cholesterol and an increased risk of atherosclerosis for the affected individuals [34]. This finding has inspired the creation of PCSK9 inhibitors as a treatment for high cholesterol, and highlights the potential for the investigation of pseudogenes to shed light on biological processes of interest to the biomedical and pharmaceutical industry [35]. However, LOF events resulting in adaptive benefits are rare.

Taken together, the well-defined evolutionary relationships between mouse strains and the wealth of associated functional data from the ENCODE project present an opportunity to investigate the processes underlying pseudogene biogenesis and activity to an extent previously not possible. Leveraging mouse developmental time-course RNA sequencing (RNA-seq) data, we explored whether pseudogene creation occurs primarily in the gametes or earlier in development in a germline precursor. In the future, comparison to the primate lineage and human population is possible as the evolutionary distance between some of the mouse strains parallels the human-chimp divergence as well as distances between the modern day human populations in terms of generations. Thus, the collection of high-quality genomes and associated pseudogene annotations for the 18 strains is a valuable resource for both population studies and the broader mouse genetics research community.

## Results

### 1. Annotation

We present the latest pseudogene annotations for the mouse reference genome as part of the GENCODE project, as well as updates on the human pseudogene reference set. Leveraging the recently assembled high-quality genome sequences of 18 mouse strains, we introduce a first draft annotation of the pseudogene complement in these genomes.

#### 1.1 Reference genome

Using a combination of rigorous manual curation [36, 37] and automatic annotation with PseudoPipe [38], we identified 20,397 pseudogenes in the mouse reference genome C57BL/6J (**Table S1A&B**). The annotation files are available to download from mouse.pseudogene.org. The present dataset is a snapshot in an ongoing annotation process. As pseudogene assignments are highly dependent on the quality of the protein-coding annotation, the current manually curated set (10,524 entries) provides a high-quality lower bound with respect to the true number of pseudogenes in the mouse genome, while the automatic annotation informs on the upper limit of the pseudogene complement size (**Figure 1C**). In agreement with our previous work [36, 37], the manual and the PseudoPipe automatic annotation show considerable overlap (over 83%) (**Table S1A**).

For human, we used a similar workflow to refine the reference pseudogene annotation to a high-quality set of 14,650 pseudogenes. The updated set contains considerable improvements in the characterization of pseudogenes of previously unknown biotype (**Table S1C**). In both the human and mouse reference genomes, the majority of the annotations are processed pseudogenes, with a smaller fraction of duplicated pseudogenes (**Table S1C**).

#### 1.2 Mouse strains

The Mouse Genome Project has sequenced and assembled genomes for 12 laboratory and four wild-derived mice, and developed a draft annotation of each organisms’ protein-coding genes [39]. Another two distant Mus species, *Mus Caroli* and *Mus Pahari*, were also sequenced and assembled [40]. Collectively, the 18 strains provide a unique overview of mouse evolution. The strains are broadly organised into three classes (**Table S2**): outgroup strains, including two independent mouse species, *Mus Caroli* and *Mus Pahari*; wild-derived inbred strains, including the subspecies *Mus Spretus* and three musculus strains (*Mus Musculus Castaneus, Mus Musculus Musculus*, and *Mus Musculus Domesticus*); and laboratory inbred strains (12 in total). A detailed summary of the genome composition for each strain is presented in [39] and [40].

We developed an annotation workflow for identifying pseudogenes in the 18 mouse strains by leveraging the in-house automatic pipeline PseudoPipe and the set of manually curated pseudogenes from the mouse reference genome lifted over onto each individual strain (**Figure 1B**). This combined pseudogene identification process gives rise to three confidence levels reflecting the annotation quality: *Level 1* includes high confidence pseudogenes with supporting evidence from both manual and automatic annotation pipelines; *Level 2* includes pseudogenes that are supported just by manual informed annotation, and *Level 3* includes pseudogenes identified only by the automatic annotation pipeline. Each identified pseudogene is associated with details about its transcript biotype, genomic location, structure, sequence disablements, and confidence level. A detailed overview of pseudogene annotation statistics including the number of pseudogenes, their confidence levels, and their biotypes is shown in **Figure 1C**.

One challenge in developing a reliable annotation is identifying the optimal trade-off between the pipeline *specificity* in producing highly accurate predictions, and *sensitivity* in estimating the total number of pseudogenes in the genome. In developing a widely used annotation resource, we want as few false positive calls as possible. Conversely, when estimating the total number of pseudogenes in a given organism, we want a more balanced ratio of false positive to false negatives. Thus, as we aimed to produce annotation files for GENCODE that would serve as the gold standard in mouse pseudogene curation, we used as input only the conserved number of protein-coding genes between each analysed strain and the reference genome. Consequently, the present workflow was fine tuned to reduce the number of false positive annotations. However, this reduces the number of pseudogenes in distant species compared to the number in the reference genome. This reduction is correlated to the decrease in the number of conserved input protein-coding genes (**Figure SF1A&B**) and informs us on the lower bound of expected pseudogene complement size.

In addition, we estimated the total number of pseudogenes in each of the 18 mouse genomes (**Table S3**) by leveraging the close relationship between the mouse reference strain C57BL/6J and its related laboratory inbred strain, the “reference-like” counterpart C57BL/6NJ (see Methods). These two strains are closely related and should have pseudogene complements of similar size. However, the strains differ in the depth of their coding transcript annotations. This enables us to estimate the impact of differential annotation depth on the number of pseudogenes identified via the present workflow.

These results suggest that all the studied strains have pseudogene complements of similar size, and that we can reduce the difference between the number of annotated pseudogenes and the expected total by improving the protein-coding annotation in each of the studied strains.

Currently, around 30% of pseudogenes in each strain are defined as high confidence Level 1 (informed from both manual curation and computational methods), 10% are Level 2 (characterised only using the lift over of the manually annotated set), and 60% are Level 3 (identified solely by the automatic annotation pipeline). The pseudogene biotype distribution across the strains closely follows the reference genome and is consistent with the biotype distributions observed in other mammalian genomes (e.g., human [36] and macaque [37]). As such, the bulk (~80%) of the annotations are processed pseudogenes, and only a smaller fraction (~15%) are duplicated pseudogenes. Finally, the density of pseudogene disablements follows the previously observed distributions in the mouse reference genome and other mammals, with stop codons being the most frequent defect per base pair followed by deletions and insertions (**Figure SF2A&B**). As expected, older pseudogenes are enriched in the number of disablements compared with the parental gene sequence. The proportion of pseudogene defects exhibits a linear inverse correlation with the pseudogene age, expressed as the sequence similarity between the pseudogene and the parent gene.

#### 1.3 Unitary pseudogenes

Unitary pseudogenes are the result of a complex interplay between LOF events and changes in evolutionary pressures that lead to the fixation of an inactive element in a species. The importance of unitary pseudogenes resides not only in their ability to mark LOF events, but also in their potential to highlight changes in the selective pressures guiding genome evolution. Due to their formation as a result of gene inactivation, identifying unitary pseudogenes is highly dependent on the quality of the reference genome protein-coding annotation, and requires a large degree of attention during the annotation process.

These pseudogenes are defined relative to the functional protein-coding elements in another species. Using a combination of multi-sequence alignments, manual curation, and a specialised unitary pseudogene annotation workflow (**Figure 1B**) in human and mouse, we identified 88 and 131 new unitary pseudogenes, respectively (**Figure 2A**). These results bring the total number of unitary pseudogenes in mouse to 165 and raise the size of unitary pseudogene class in human to 303 entries (**Table S4**). This is a considerable increase compared to previous GENCODE releases [21, 41, 42] and can be largely attributed to improvements in mouse genome annotation and assembly. In human, a large proportion of the new unitary pseudogenes are related to the chemosensory system (e.g. olfactory receptor proteins); these pseudogenes have functional homologs in mouse, reflecting the LOF in these genes during the human lineage evolution. A known example of human unitary pseudogene which has functional counterparts in mouse, rabbit, and several primates is with respect to the Cyp2G1 mouse protein (**Figure 2B**). Here, the human gene acquired a C-to-T mutation equating to a stop codon in the middle of a coding exon, which resulted in gene disablement and thus the creation of a unitary pseudogene.

**Figure 2.**
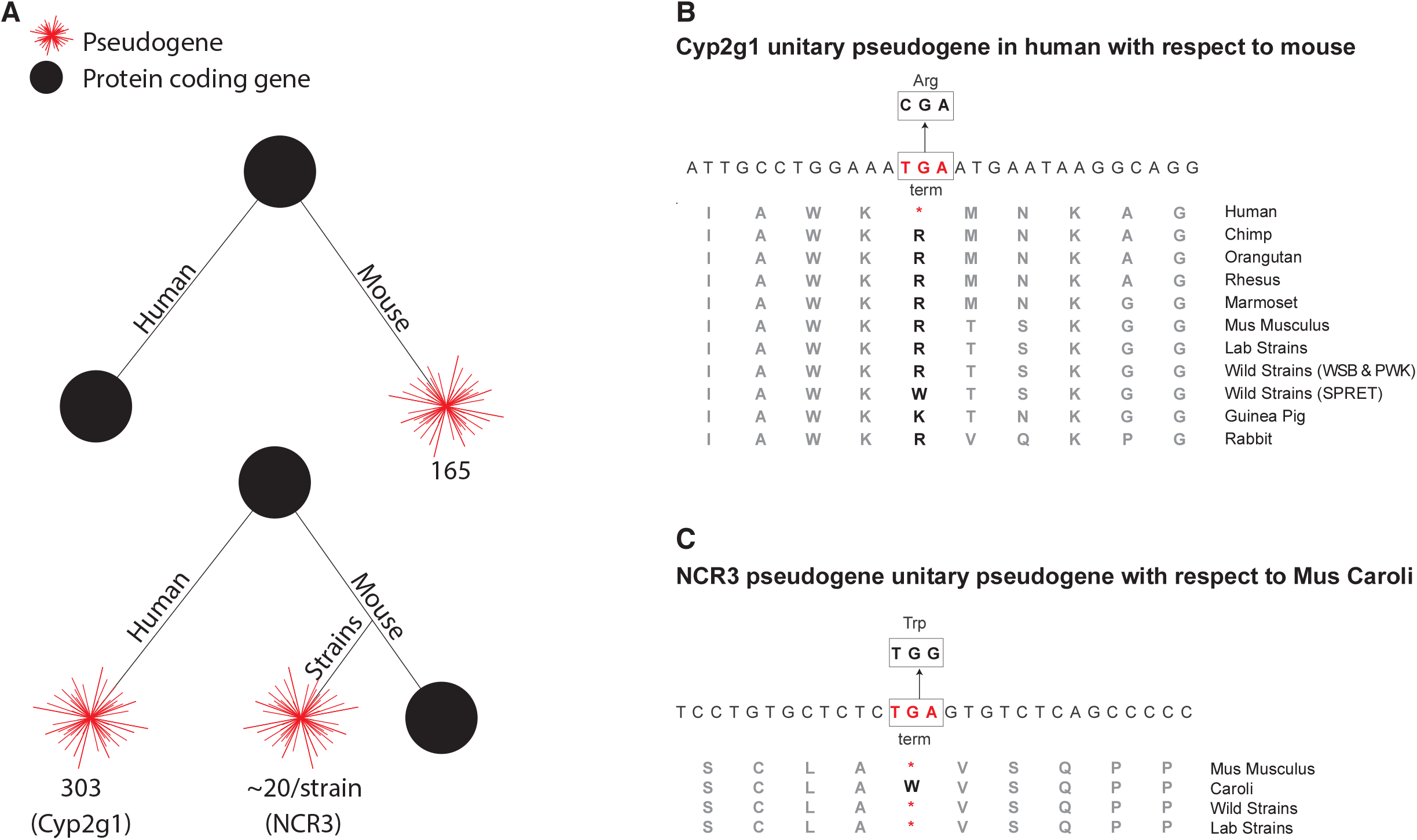
A – Summary of unitary pseudogenes with respect to human and mouse. B – Cyp2G1 LOF in human. C – NCR3 GOF mutation in *Mus Caroli* as compared to the reference genome and the other mouse strains.

Moreover, we observed the pseudogenisation of a number of innate immune response related genes in humans such as Toll-like receptor gene 11 and leucine rich repeat protein gene, hinting at potentially advantageous LOF/pseudogenisation events in human lineage evolution [43]. In mouse, the majority of the new unitary pseudogenes with respect to human are associated with structural Zinc finger domains and immunoglobulin proteins (**Table S5**).

Focusing on the functional paralogs, we observed that new unitary pseudogenes are in general associated with lowly expressed protein-coding genes in their corresponding organisms. In particular, more than half of the human functional orthologs that have been pseudogenised in mouse show no level of expression in the top-tier ENCODE cell lines (**Figure SF3A**). The top broadly expressed protein-coding genes are Ribosomal proteins, NADH ubiquinone oxidoreductase, and the solute carrier SLC25A6. Additionally, the majority of these functional genes are classified as non-essential according to the OGEE database, suggesting that the null deletions of these genes are less likely to have a negative impact on the organisms’ survival.

Similarly, a large fraction of the mouse functional orthologs of the human unitary pseudogenes show low levels of expression across 18 mouse tissues (**Figure SF3B**). Moreover, those that are transcribed are tissue specific.

The draft nature of the mouse strains’ annotation and assembly makes it difficult to identify unitary pseudogenes within them. To get an overview of the unitary pseudogenes in each strain, we used the mouse reference genome as the required canonical organism and followed a similar workflow as described above. However, in this case we used mouse reference peptides that are not present in the analysed strain, as input. We intersected the resulting pseudogene calls with the lift over of protein-coding genes specific to the reference in order to validate the conservation of location and LOF of the reference protein coding genes in the analysed strain. On average, we found around 20 unitary pseudogenes in each strain, with larger numbers for wild-derived inbred strains (**Table S6**). We can distinguish the unitary pseudogenes from other strain-specific pseudogenes by the fact that they do not have a functional homolog (parent gene) in the same organism. Moreover, the fast rate of evolution among the mouse strains, as well as the highly specific generation of the laboratory strains, suggests that the number of unitary pseudogenes could be considerably higher, reflecting the strain-specific phenotypes. A way to realistically assess the size of the unitary pseudogene complement is to look at the unitary annotation in the human genome relative to other primates [21], as previous studies suggest that the protein loss rate is comparable in both rodent and primate lineages [42]. As such, we expect that the total number of unitary pseudogenes with respect to the reference in each strain will depend on the evolutionary distance between the two and will be comparable to the number of human-specific unitary pseudogenes with functional homologs in chimp (estimated to be 403 [44]).

Similarly, future improvements in the strain annotation will allow us to annotate unitary pseudogenes in the reference with respect to the mouse strains. These elements will highlight not only LOF events in the reference, but also fixation of GOF mutations in divergent strains and species. we found that this is the case for the NCR3 gene in *Mus Caroli* (**Figure 2C**). Here, we observed an A-to-G GOF mutation for the NCR3 gene that is pseudogenised in all the other mouse strains including the reference, reverting the initial TGA stop to a tryptophan codon.

### 2. Conservation and divergence in pseudogene complements

In order to investigate the evolutionary history of pseudogenes in the mouse strains, we created a *pangenome* pseudogene dataset containing 49,262 unique entries relating the pseudogenes across strains. We found 2,925 pseudogenes that are preserved across all strains. A detailed summary of the other subsets of pseudogenes is shown in **Figure 3A,B**. On average, each strain contains between 1,000 and 3,000 pseudogenes that are not directly associated with any pseudogenes in the other strains, based on our imposed ortholog selection criteria (see **Methods**). By relaxing these constraints, we were able to estimate the minimum number of strain-specific pseudogenes. To this end, we identified on average 293 unique elements in each analysed genome. However, this is only a lower bound estimate. Moreover, the proportion of pseudogenes conserved only in the outgroup, the wild-derived strains, or the lab strains is considerably smaller, suggesting that the bulk of the pseudogenes in each strain was created during the shared evolutionary history. Additionally, the largest group of pseudogenes is composed of those with orthologs in at least one other strain. As the classical laboratory strains have been derived mainly from WSB (94.3± 2.0%), with smaller contribution from PWK (5.4± 1.9%) and CAST (0.3± 0.1%) [45], we expect a very small fraction of the identified orthologs to have no evolutionary significance.

**Figure 3.**
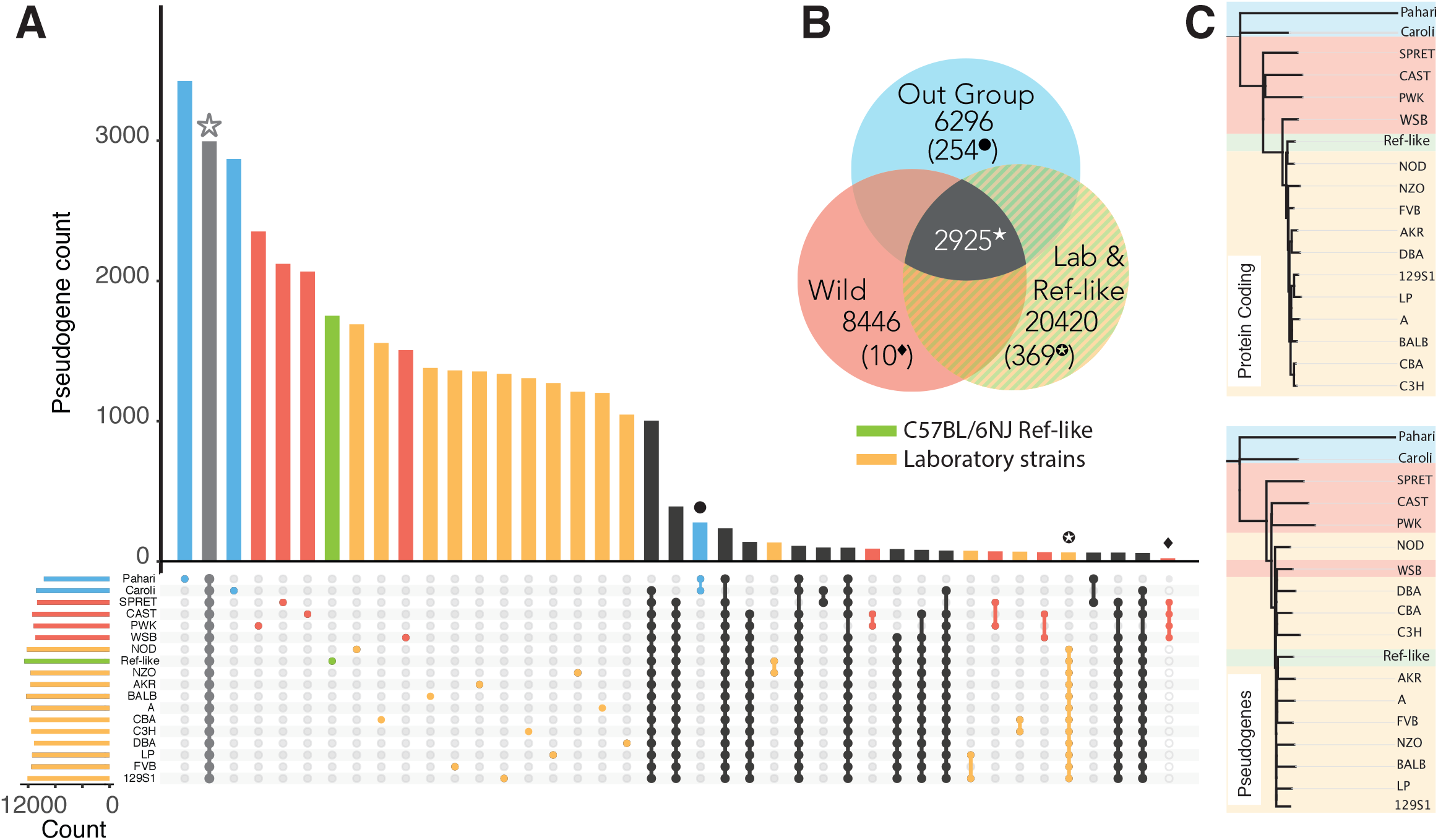
A – Summary of pseudogene distribution in the pangenome mouse strain dataset. The different group of mouse strains are highlighted by colours: blue relates to outgroup mice (*Mus Pahari* and *Mus Caroli*), red corresponds to wild-derived mice (SPRET, CAST, WSB, PWK), yellow indicates the laboratory inbred strains as listed in **Table S2**, and green highlights the laboratory inbred “reference-like” strain C57BL/6NJ. B – Venn diagram of evolutionarily conserved and group-specific pseudogenes. The number in brackets is indicative of pseudogenes that are unique to each group. C – Phylogenetic trees for parents of evolutionarily conserved pseudogenes and evolutionary conserved pseudogenes. Bootstrap values are provided in mirror figure **SF3D**.

The evolutionary information recorded in the pangenome dataset allowed us to infer a pseudogene’s age based on its presence or absence in syntenic regions across various strains. To this end, we constructed a binary matrix from the pangenome dataset and estimated the relative age of the pseudogenes as the age of strain divergence (**Figure SF3C**). This approach however, does not account for pseudogenes that have been lost in various strains after speciation. Furthermore, clustering the strains based on the absence or presence of pseudogenes in syntenic regions recovers the known speciation events in the mouse lineage showcasing pseudogenes as markers of genome evolution.

Next, we took advantage of the ability of pseudogenes to evolve with little or no selective constraints [46], and compared sequence divergence across the mouse strains. To this end, we built a phylogenetic tree based on approximately 1,500 pseudogenes that were randomly selected and are conserved across all strains (**Figure 3C, SF3D**). This pseudogene-based tree follows closely the tree constructed from protein-coding genes and correctly identifies and clusters the mice into three classes: outgroup, wild-derived, and laboratory strains. In constructing these trees, we concatenated the gene sequences in the same order in which they occur in the various strains. The use of pseudogenes and protein-coding genes across the genome reduces any potential bias introduced by the strains’ mosaicism by averaging out the contribution of any given genomic region. Additionally, the use of pseudogenes conserved across all strains removes any pseudogenes shared among subsets of strains due to contamination via introgression.

### 3. Genome Evolution and Plasticity

Leveraging the pseudogene annotations, we explored the differences between the mouse strains by looking at the genome remodeling processes that shaped the evolutionary history of their pseudogene complements.

#### 3.1 Pseudogene Genesis

Taking advantage of the available functional genomics and evolutionary data, we can study pseudogene genesis on a unique scale: during embryo development at one extreme and the mouse lineage at the other.

Given that processed pseudogenes are formed through the retrotransposition of the parent mRNAs, it is reasonable that the expression level of the parent gene directly correlates with the number of processed pseudogenes [47]. Moreover, as pseudogenes are inherited, the genesis of new elements occurs in the germline. we used an embryogenesis RNA-seq time course dataset to test our assumptions during early development [48]. We calculated the parent gene expression for a series of developmental stages ranging from metaphase II oocytes to the inner cell mass. At every stage, the average expression level of parent genes is higher than that observed for non-parent protein-coding genes. However, genes associated with large pseudogene families show low transcription levels during very early development, with high expression levels achieved only during later stages. We evaluated the correlation between the expression level of a gene and the number of pseudogenes associated with it at different developmental timepoints. This correlation improves as we progress through the developmental stages, suggesting that pseudogenes are most likely generated by highly expressed housekeeping genes (**Figure SF4A,B**). Next, we explored in a similar fashion the parent gene expression during spermatogenesis using data from Hammoud *et al.* (2014) [49]. We did not find any strong trends between parent gene expression and the number of pseudogenes, regardless of the male germ line developmental stage (**Figure SF4C**).

We further tested the correlation between high expression levels and the number of associated pseudogenes by looking at RNA-seq samples from adult mouse brain. Similar to our previous observations, the pseudogene parent genes show a statistically significant increase in average expression levels compared to non-pseudogene-generating protein-coding genes (**Figure SF4D,E**).

Next, we examined the degree to which the number of pseudogenes is related to the number of copies or functional paralogs of the parent gene (**Figure 4A**). For duplicated pseudogenes, we observed a weak correlation between the number of paralogs and the number of pseudogenes of a particular parent gene. This result suggests that a highly duplicated protein family will tend to give rise to more disabled copies than a less duplicated family, if we assume that each duplication process can potentially give rise to either a pseudogene or a functional gene. This distribution of duplicated pseudogenes to duplicated paralogs in mouse is in accordance with previous observations in human [37, 50].

**Figure 4.**
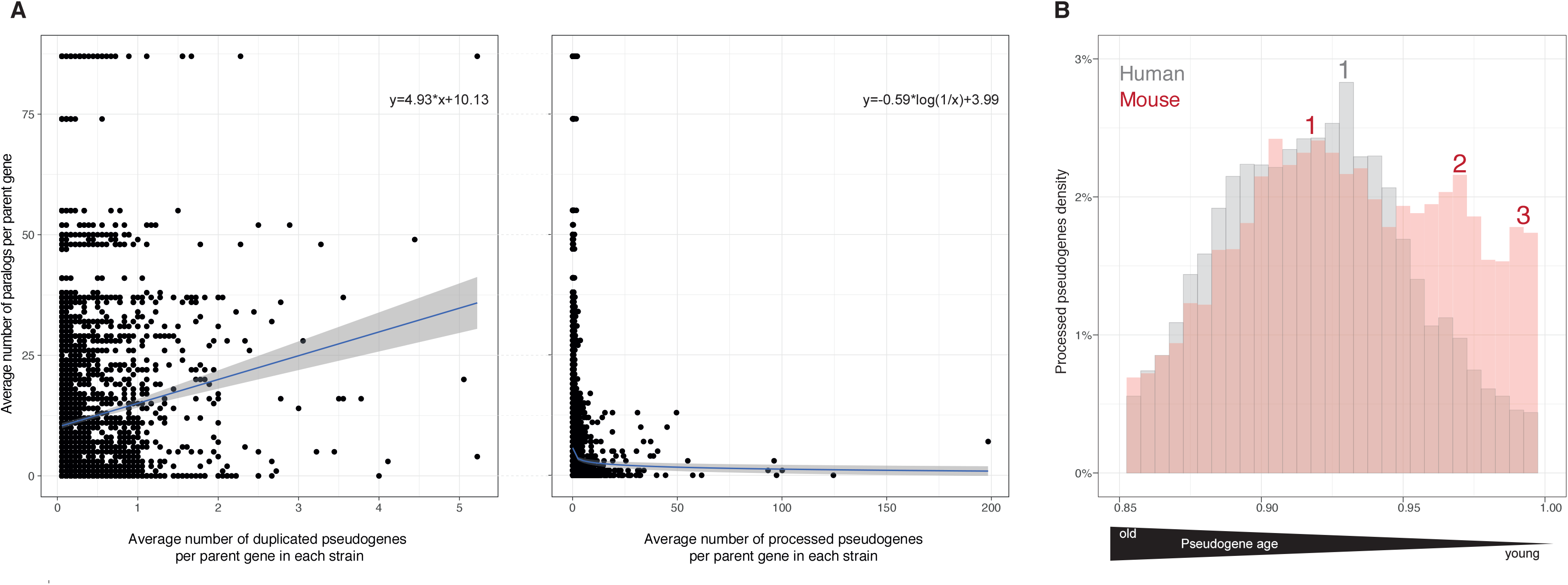
A – Relationship between the number of pseudogenes and functional paralogs for a given parent gene (left – duplicated pseudogenes, right – processed pseudogenes). Fitting lines show a vague correlation between the number of functional vs. disabled copies of a gene, with a linear fit for duplicated pseudogenes and a negative logarithmic fit for processed pseudogenes. The gray area is the standard deviation. B – Distribution of processed pseudogenes (y-axis) as a function of age (x-axis). The pseudogene age is approximated as DNA sequence similarity to the parent gene.

By contrast, for processed pseudogenes we observed a weak inverse correlation. This result implies that in the case of large protein families we can expect to see a lower level of transcription for each family member, with high mRNA abundance being achieved from multiple duplicated copies of a gene rather than increased expression of a single unit. This is particularly the case for parent genes from the large PRAME (preferentially expressed antigen in melanoma) family, which have more than 80 paralogs in the mouse genome and less than two or even no processed pseudogenes. These genes showed low or no expression in the studied adult brain (FPKM~0). On the opposite side of the spectrum, the GAPDH gene family, which has only seven paralogs in mouse and almost 200 processed pseudogenes, showed a significantly higher expression level (FPKM>8) in the adult mouse brain (**Figure SF5A**).

An interesting question on the relationship between the number of paralogs in a given gene family and the number of associated pseudogenes is whether this relationship is driven by the family of genes (e.g., olfactory receptors for duplicated pseudogenes and ribosomal proteins for processed pseudogenes). To answer this question, we restricted our analysis to these two families (**Figure SF5B**). However, we did not find any family-specific trends. This might be due to the low number of data points compared to the full dataset.

#### 3.2 Transposable elements

As the majority of mouse and human pseudogenes are the result of retrotransposition processes mediated by transposable elements (TEs), we investigated the genomic mobile element content in human and mouse as well as the generation of processed pseudogenes on an evolutionary time scale (**Figure 4B, SF5C**).

TEs are sequences of DNA characterised by their ability to integrate themselves at new loci within the genome. TEs are commonly classified into two classes: DNA transposons and retrotransposons, with the latter being responsible for the formation of processed pseudogenes and retrogenes. Both human and mouse genomes are dominated by three types of TEs, namely short interspersed nuclear elements, long interspersed nuclear elements (LINEs), and the endogenous retrovirus superfamily. LINE-1 (L1) elements have been shown to mobilize Alus, small nuclear RNAs, and mRNA transcripts. We analysed the L1 retroposed pseudogenes in human and mouse. We defined the evolutionary time scale by using the pseudogene sequence similarity to the parent gene as a proxy for age. Younger pseudogenes have a higher degree of sequence similarity to the parent, whereas older pseudogenes show a more diverged sequence.

In human, we observed a smooth distribution of processed pseudogenes, with a single peak at 92.5% nucleotide sequence similarity to parents. This finding hints at a burst of retrotransposition events that occurred 40 MYA at the dawn of primate lineage and created the majority of human pseudogene content [51, 52]. By contrast, in mouse we found that the processed pseudogene distribution is defined by two successive peaks at 92.5% and 97% sequence similarity to parent genes. Also in contrast to human where the density of processed pseudogenes shows a steep decrease among young pseudogenes following the peak at 92.5% similarity, the density of mouse processed pseudogenes remains high in the interval from 97% to 100% sequence similarity to parents. A close examination of the young pseudogene age distribution suggests a reduction in the number of newly created pseudogenes. This is most likely a consequence of the stringent criteria used in calling pseudogenes at high sequence similarity to parents and showcases the difficulty in annotating recently disabled/dead genes due to their high similarity to functional protein-coding counterparts. Overall, these results suggest the presence of active transposable elements in mouse, which results in continuous renewal of the processed pseudogene pool. This is also reflected in the large difference in the number of active LINE/L1s between human and mouse, with just over 100 in human [53] compared to 3,000 in mouse [54].

#### 3.3 Genome remodeling

The large proportion of strain- and class-specific pseudogenes, as well as the presence of active TE families, point towards multiple genomic rearrangements in mouse genome evolution. To this end, we examined the preservation of pseudogene genomic loci between each of the mouse strains and the reference genome for one-to-one pseudogene orthologs (**Figure 5A,B**). We observed that on average more than 97.7% of loci are preserved across the laboratory strains, and 96.7% of loci are preserved with respect to the wild-derived strains. By contrast, only 87% of *Mus Caroli* loci and 10% of *Mus Pahari* loci are preserved in the reference genome. The significant drop in the number of preserved pseudogene loci between the reference genome and outgroup strains is in agreement with the observed major karyotype-scale differences and large genomic rearrangements exhibited by *Mus Caroli* and *Mus Pahari* [40, 55]. The proportion of non-preserved loci follows a logarithmic curve that matches closely the divergent evolutionary time scale of the mouse strains (**Figure 5C**). This suggests a uniform rate of genome remodeling processes across the murine taxa. Next, we examined the distribution of preserved and non-preserved loci as a function of strain divergence times for processed and duplicated pseudogenes (**Figure SF6**). While overall the pseudogenes seem to follow the same logarithmic curve, we noticed that the deviation from the perfect fit is larger for non-processed pseudogenes compared to processed ones. This difference, however, can be attributed to the considerable differences in the input dataset size.

**Figure 5.**
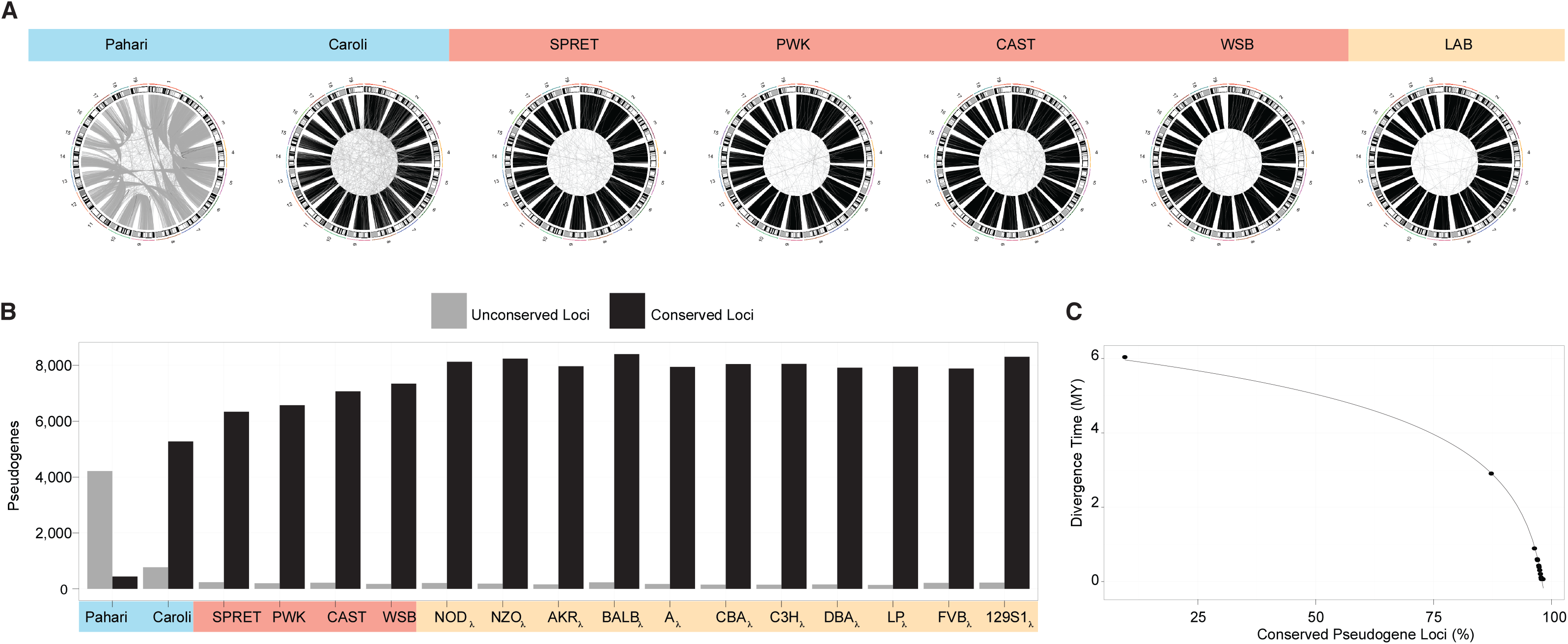
A – CIRCOS-like plots showing the conservation of the pseudogene genomic loci between each mouse strain and the laboratory reference strain C57BL/6NJ. Gray lines indicate a change of the genomic locus between the two strains and connect two different genomic locations (e.g., a pseudogene located on chr7 in C57BL/6NJ and chr1 in *Mus Pahari*). Black lines indicate the conservation of the pseudogene locus. B – The number of pseudogenes that are preserved or have changed their loci between each strain and the laboratory reference strain. B – The number of pseudogenes that are preserved or changed their loci between each strain and the laboratory reference strain. C – Strain speciation times as a function of percentage of conserved pseudogene loci between each strain and the laboratory reference, fitted by an inverse logarithmic curve.

### 4. Functional analysis

The role of pseudogenes in genome biology has long been debated. However, recent studies [37] have highlighted the fact the pseudogenes can reflect the evolution of genome function and activity. Here, we addressed the biological relevance of pseudogene activity by leveraging data from gene ontology (GO), protein families, and RNA-seq experiments.

#### 4.1 Gene ontology and pseudogene family analysis

We integrated the annotations with GO data in order to characterise the functions associated with pseudogene generation. For this, we calculated the enrichment of GO terms across the strains. We observed that the majority of top biological processes, molecular functions, and cellular component GO terms are shared across the strains (**Figure 6A**). We also evaluated GO term enrichment among parent genes for both processed and duplicated pseudogenes across the mouse strains. Enriched GO terms were clustered based on semantic similarity and the strains were clustered based on GO term enrichment profile similarity. The resultant heatmap (**Figure 6B**) enables us to identify both related terms with conserved enrichment across all strains as well as blocks of terms that exhibit conservation within a single or a few closely related strains. We observed Conserved enrichment for GO terms related to ribosomal functions, cell cycle, translation and RNA processing, and ubiquitination for processed pseudogenes. Among duplicated pseudogenes, we observed enrichment for apoptosis, sensory and smell processes, and immune functions. Additionally, the GO terms that universally characterise the pseudogene complements in all the mouse strains are closely related to the family classification of pseudogenes. As we reported previously, the top pseudogene family is 7-Transmembrane [37]. This Pfam family encompasses the chemoreceptors’ GPCR proteins, reflecting the enrichment in olfactory receptors in the mouse. Similar to the human and primate counterparts, many top families in mouse pseudogenes are related to highly expressed and duplicated proteins such as GAPDH and ribosomal proteins, and regulatory protein families such as the Zinc fingers (**Figure 6C**) [56].

**Figure 6.**
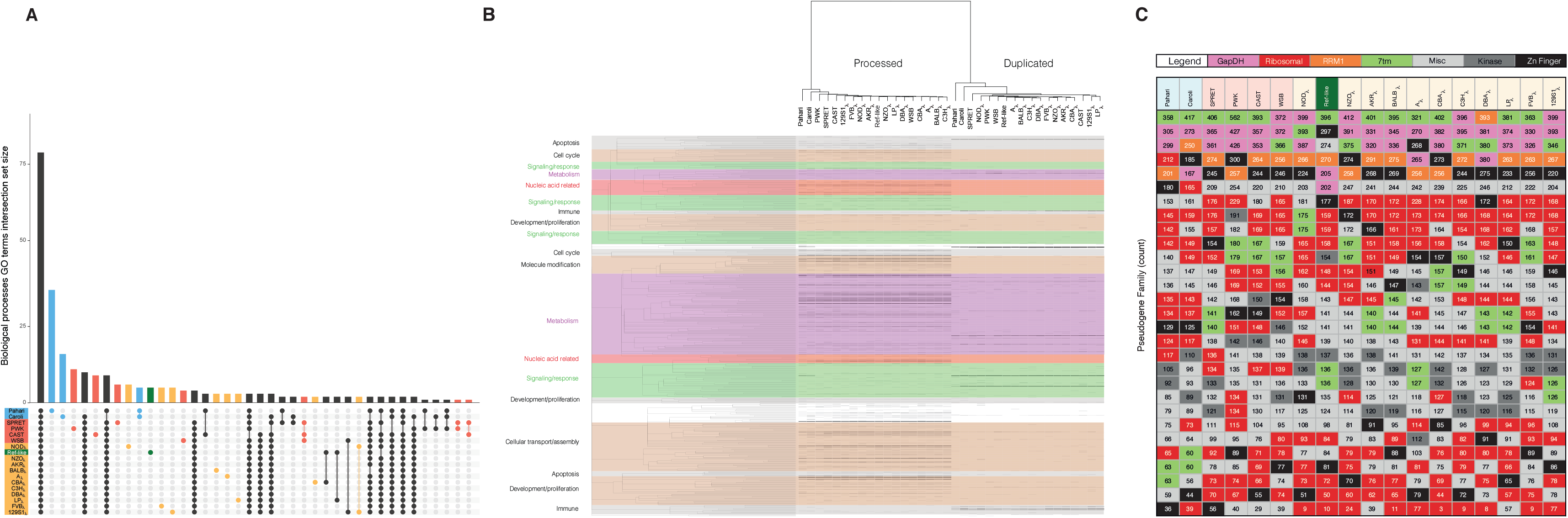
A – Distribution of enriched GO biological processes terms across the mouse strains. B – Heatmap illustrating enrichment of GO biological processes terms across the mouse strains for the parent genes of processed and duplicated pseudogenes. GO terms (rows) are clustered by semantic similarity (colour). Each line in the heat map indicates the presence of a pseudogene. The GO terms shown in colour indicate an association with the pseudogene family of similar colour in panel C. C – Summary of the top 24 Pfam pseudogene families in each mouse strain.

A closer look suggests that the pseudogene repertoire also reflects individual strain-specific phenotypes (**Table S5**). We can view the pseudogene-phenotype relationship from different perspectives. First, pseudogenes reflect duplication events linked with the emergence of an advantageous phenotype. This is the case for the *Mus Spretus* genome, where we observed an enrichment of duplicated tumor repressor and apoptosis pathways genes [57] and a corresponding increase in the number of associated pseudogenes. Second, we found pseudogenes reflecting the death of a gene family. As such, we observed an increase in the number of pseudogene-associated deleterious phenotypes. A well-known example is the pseudogenization of Cytochrome c Oxidase subunit VIa through accumulation of LOF mutations in the blind albino mouse strain, which is commonly linked with neurodegeneration [58] and is characteristic for the observed brain lesions in the affected mice [14]. However, a detailed analysis of the pseudogene repertoire suggests that there are more ways to describe the pseudogene–phenotype association, in particular by considering the emergence of advantageous phenotypes through the pseudogenization process [59].

#### 4.2 Gene essentiality

We observed an enrichment of essential genes among pseudogene parent genes in the mouse strains. Evaluating the parent gene for each pseudogene present in the mouse strains revealed that essential genes are approximately three times more abundant among parent genes (**Table S7A**). In general, essential genes are more highly transcribed than non-essential genes [60], and thus might be associated with a higher propensity of generating processed pseudogenes. However, one potential confounder is the gene expression level, which is associated with genes that are more processed or more essential. Thus, we evaluated the probability that a gene is essential by controlling for its transcription level and parent gene status (see **Methods**), and found that pseudogene parents are still 20% more likely to be essential genes compared to regular protein-coding genes (**Table S7B**).

We also analysed the number of paralogs associated with our essential and non-essential gene sets to gain insight into the possible role of gene duplication in the enrichment of essential genes among the parent genes set. In the reference mouse, 80.6% of non-essential genes and 74.1% of essential genes have paralogs. This is in agreement with previous work showing that non-essential genes are more likely than essential genes to be duplicated successfully [61].

#### 4.3 Pseudogene Transcription

We leveraged RNA-seq data from the Mouse Genome Project and ENCODE to study pseudogene biology as reflected by their transcriptional activity. This is thought to either relate to the exaptive functionality of pseudogenes or be a residual leftover from their existence as genes. In both the human and mouse reference genomes, we found that about 15% of pseudogenes were transcribed across a variety of tissues, a result similar to previous pan-tissue analyses (**Figure 7A,B**) [36, 37].

Due to restricted data availability in the mouse strains, we focused our transcriptional analysis to a single tissue: adult brain from wild-derived and laboratory strains. Overall, pseudogenes with strain-specific transcription were more common than those with cross-strain transcription (**Figure 7C,D**). Moreover, the proportion of pseudogenes conserved across all strains that are transcribed is constant (~2.5%) across the wild-derived and laboratory strains (**Figure 7D**). By contrast, the fraction of transcribed strain-specific pseudogenes varies across the strains from 1.5% to 4% (**Figure 7D**).

**Figure 7.**
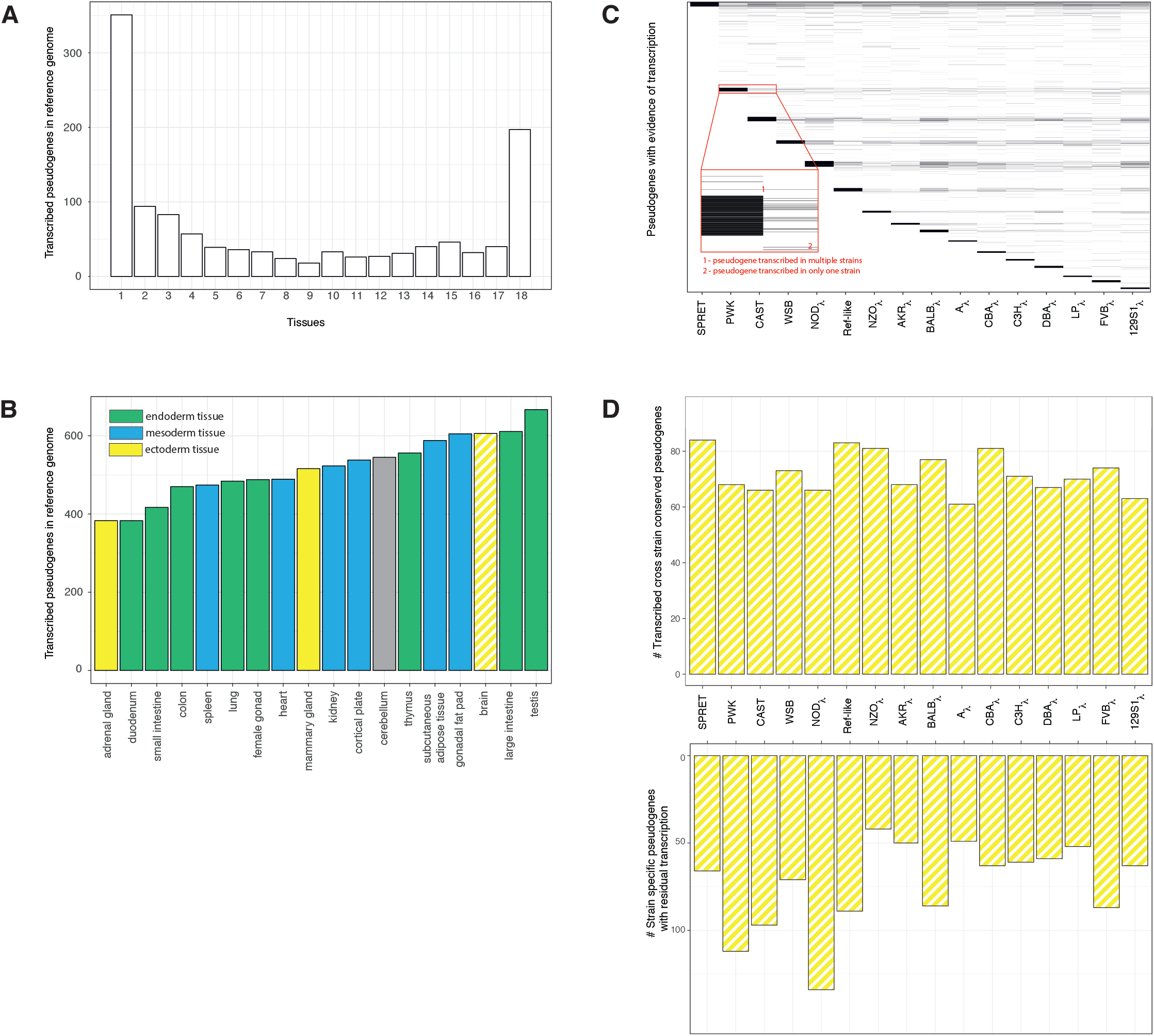
A – Cross-tissue pseudogene transcription in the mouse reference genome. The x-axis indicates the number of tissues in which a pseudogene is transcribed. B – Distribution of pseudogene transcription in 18 adult mouse tissues. C – Heatmap-like plot showing the distribution of transcribed pseudogenes (y-axis) in brain tissue for each wild-derived and laboratory mouse strain (x-axis). Each line corresponds to a transcribed pseudogene with an expression level higher than 2 (FPKM). When a line is present across multiple columns, it is indicative of a pseudogene expressed in all these strains. The dark bars at the top of each strain column are formed by multiple highly expressed pseudogenes. When a line is present in only one strain, and no other line is observed at the same level in any of the other strains, this suggest that the pseudogene expression is strain specific. D (top) – number of transcribed pseudogenes that are conserved across all the strains. D (bottom) – number of transcribed strain-specific pseudogenes in each mouse strain.

### 5. Mouse pseudogene resource

We created a comprehensive resource that organises all of the pseudogenes across the available mouse strains and the reference genome, as well as associated phenotypic information. This database is available online at mouse.pseudogenes.org. The database contains information regarding strain and cross-strain annotation, pseudogene family and phenotypic data, as well as expression data. All the available data are provided as flat files for ease of manipulation.

Queries on specific pseudogenes will return the relevant pseudogene annotation containing all pertinent associated information. The pseudogenes are annotated with a unique universal identifier as well as a strain-specific ID in order to facilitate both the comparison of specific pseudogenes across strains and collective differences in pseudogene content between strains. This enables pairwise comparisons of pseudogenes between the various mouse strains and the investigation of differences between multiple strains of interest.

## Discussion

In this study, we report the updated and refined pseudogene annotation in the mouse and human reference genomes as part of the GENCODE genome annotation project and describe the curation and comparative analysis of the first draft of pseudogene complements in 18 related mouse strains. By combining computational and manually informed annotations, we obtained a comprehensive view of the pseudogene content in genomes throughout the mouse lineage. The overlap between manually curated pseudogenes and those identified using Pseudopipe (in-house computational method) is over 80%, reflecting the high sensitivity of the computational detection method.

The reported number of pseudogenes for each of the mouse strains in our annotation files is a conservative estimate of the total in order to down weight the number of potential false positives. Also, it represents a snapshot in an ongoing annotation process. Using information from manual curation and automatic annotation, we also computed an estimate of the total pseudogene complement size in each of the strains with a more balanced ratio of false positives to false negatives. Our previous experience with human genome annotation suggests that improvements in manual curation translate to a higher-quality pseudogene dataset [36, 37]. Specifically, we expect the proportion of high confidence level pseudogene annotations (Level 1) to increase, consequently leading to a decrease in the number of pseudogenes identified only by computational methods (Level 3). we plan to update all of the annotations in line with major reference genome releases. Furthermore, any updates will ensure consistency in the nomenclature of pseudogene biotypes. This is particularly the case for polymorphic and transcribed pseudogenes. Potentially, these could be later classified as protein-coding genes subjected to LOF disablements in various individuals within a population and long non-coding RNA (ncRNA), respectively.

Comparable to our previous observations in human, worm, and fly [37], the pseudogene complement in mouse strains reflects an organism-specific evolution. Despite this, the pseudogenes share a number of similarities regarding their biogenesis and diversity. As such, we noticed a uniform ratio of processed-to-duplicated pseudogene of 4 to 1 in all of the strains, a result consistent with previous observations in human [36, 37]. The higher proportion of processed pseudogenes, accounting for ~80% of the total, is in agreement with earlier findings that suggest that retrotransposition is the primary mechanism for pseudogene creation in numerous mammalian species [36]. Moreover, when we examined the retrotransposon activity, and in particular the L1 content, we observed that while the majority of human pseudogenes have been formed relatively recently through a single burst of retrotransposition [36], the mouse lineage shows sustained renewal of the pseudogene pool through successive bursts. The sequence context of the processed pseudogenes indicates that the various retrotransposons exhibit differential contributions to the pseudogene set over time.

As a pseudogene’s likelihood of creation is related to its parent gene’s functional role and expression level, it can act as a record of its parent’s past importance and expression level. The link between the creation of processed pseudogenes and parent genes associated with key biological functions is further supported by an enrichment of parent genes among essential mouse genes. Meanwhile, duplicated pseudogenes record events that shaped both the genome environment and function during an organism’s evolution. Furthermore, the wealth of functional genomics assays available for experimentally relevant mouse strains presents an opportunity to investigate both the activity of parent genes as well as pseudogene genesis. As expected, we observed that parent genes have higher levels of expression relative to non-parents during both embryonic development and adulthood. Moreover, although time series expression analyses during embryonic development did not identify a single developmental time-point at which parent gene expression was strongly associated with pseudogenesis, a clear relationship was observed in the most mature cell types, suggesting that most pseudogene creation is commonly related to the high expression levels of ubiquitous housekeeping genes.

To better understand the evolutionary and functional relationship between the pseudogenes in the 18 strains, we constructed a pan-genome pseudogene set as the union of all individual strain complements, resulting in over 45,000 unique entries. The pan-genome pseudogene repertoire distinguishes three types of pseudogenes: universally conserved (present in all 18 strains), multi-strain (present in at least two strains), and strain specific (unique to a specific strain and without an associated ortholog in another strain), accounting for 6%, 23%, and 71% of the elements, respectively. Despite the large number of pseudogenes without an associated ortholog in the pangenome set, these account for only 20% of the total reported pseudogenes in any particular strain, a comparable proportion to the universally conserved pseudogenes present in each strain. Moreover, the pseudogene cross-strain relationship gives us a closer look at the evolution of pseudogenes by studying the conservation of their chromosomal location. In particular, we observed a stark contrast between the high level of genomic loci retention shared by the laboratory strains and the lack of conservation among the outgroup species. These results hint at multiple large-scale genomic rearrangements in the mouse lineage. This was especially noticeable in *Mus Pahari*, as was recently reported in a large-scale chromosomal imagining and karyotype analysis [40, 55].

Analysis of pseudogenes and their parent genes can elucidate changing functional constraints and selective pressures. Unitary pseudogenes are markers of LOF mutations that that have become fixed in the population. Here, we annotated over 200 new unitary pseudogenes in mouse and a similar number in human. We found that the enrichment of vomeronasal receptor unitary pseudogenes in human with respect to mouse highlights the loss of certain olfactory functions in humans. Unitary analysis is especially interesting because it provides us with key moments in the evolution of gene function by marking LOF and GOF events. In particular, the human Cyp2G1 gene was pseudogenised while its mouse counterpart is still functional. Similarly, a disablement reversal mutation leads to a functional NCR3 gene in *Mus Caroli* while the mouse reference, laboratory, and wild-derived strains show the presence of a pseudogene on the same locus.

Taking advantage of information-rich resources such as GO and Pfam, we functionally characterised the pseudogenes. For this, we annotated pseudogenes and parent genes in each strain with GO terms and Pfam families. We observed an enrichment in housekeeping functions associated with conserved pseudogenes as illustrated by the presence of GAPDH, ribosomal proteins, and zinc finger nucleases as top Pfam families among the mouse pseudogenes. The top mouse pseudogene families closely match those seen in human. The GO enrichment analysis supports the above results, with top terms including RNA processing and metabolic processes. Additionally, we used the pan-genome pseudogene set to identify strain-specific functional annotations and suggest hypotheses as to what cellular processes and genes might underpin phenotypic differences between the mouse strains. For example, we observed that PWK is associated with strain-specific GO terms for melanocyte-stimulating hormone receptor activity and melanoblast proliferation, which may play a role in the strain’s patchwork coat colour [62]. NZO, an obesity prone mouse strain, was characterised by a specific enrichment in defensin-associated pseudogenes. Defensins are small peptides that control the inflammation resulting from metabolic abnormalities in obesity and type 2 diabetes [63], and were recently identified as potential markers of obesity [64]. Taken together the functional analysis of pseudogenes provides an opportunity to better understand the selective pressures that have shaped an organism’s genomic content and phenotype.

When we examined pseudogene expression across the strains, we observed evidence of pseudogenes with broadly conserved transcription as well as some with strain-specific expression. As additional RNA-seq datasets for multiple tissues for each strain become available, future work can investigate both pan-strain and pan-tissue expression patterns.

Finally, throughout our analysis, we present pseudogenes as ideal markers of genome remodeling because, in contrast to protein-coding genes, pseudogenes evolve with little or no selection pressure, are free to acquire mutations and thus provide a detailed history of a genome’s evolution. In particular, we observed that in contrast to human, mouse lineage is marked by multiple recent bursts of retrontransposition events. Furthermore, the lack of pseudogene loci conservation between outgroup species and the reference hints at large scale genomic rearrangements in the mouse lineage, while the presence and absence of pseudogenes in various strains is sufficient to inform the correct phylogeny of the three main groups of mouse strains analysed.

In summary, this comprehensive annotation and analysis of pseudogenes across 18 mouse strains provides support for conserved aspects of pseudogene biogenesis and expands our understanding of pseudogene evolution and activity. Integration of the pseudogene annotations with existing knowledge bases including Pfam and GO provided insight into the biological functions associated with pseudogenes and their parent genes. Furthermore, the well-defined relationships between the strains aided our evolutionary analysis of the pseudogene complements. Taken together, annotation of pseudogenes across a range of extensively used laboratory mouse strains and their integration into a comprehensive database with evolutionary and functional genomics data provided a useful resource for the broader research community. Additionally, the experimental and functional genomics datasets associated with these well-studied strains shed light on the transcriptional activity of pseudogenes and offer promise for future studies.

## Materials and Methods

### Code and data availability

The pseudogene annotation pipeline is freely available at http://pseudogene.org/pseudopipe. All supplementary data is available at http://mouse.pseudogene.org/Supplement/.

### Datasets

Mouse reference genome is based on the *Mus Musculus* strain C57BL/6J. The mouse reference annotation is based on GENCODE vM12/Ensembl 87.

The human reference genome annotation is based on GENCODE v25/Ensembl 87.

The 16 laboratory and wild-derived inbred strains’ (**Table S2**) assemblies and strain-specific annotations were obtained from the Mouse Genome Project [39] (http://www.sanger.ac.uk/science/data/mouse-genomes-project, last accessed on 21.08.2017). The laboratory strain C57BL/6NJ is a subline of the reference strain [14]. There is high sequence and evolutionary similarity between the reference genome single inbred strain C57BL/6J and the laboratory inbred mouse strain C57BL/6NJ. For the purpose of this study and in order to facilitate a reliable comparison across all the studied mouse genomes, we used the laboratory inbred strain C57BL/6NJ (labeled “reference-like” or “ref-like”) as a reference point.

The two outgroup mouse species (**Table S2**), *Mus Caroli* and *Mus Pahari*, were sequenced, assembled, and annotated in the protein-coding domain by [40].

### Lineage Comparisons

Human – primate lineage divergence and generation times were obtained from [65]. The divergence times for the wild-derived and laboratory strains were obtained from [66, 67, 68]. The data for the two outgroup species’ divergence times was obtained from [40]. The generation time for all the mice was estimated from [14].

### Pseudogene Annotation

#### Reference genome annotation

We manually curated 10,524 pseudogenes in the mouse reference genome (GENCODE M12) and 14,650 pseudogenes in the human reference genome (GENCODE v25), using a workflow previously described in [36, 37]. The manual annotation is based on the sequence homology to protein data from UniProt database [37] and the protocol is summarized in **Figure SF7**.

The number of manually annotated pseudogenes in the mouse lineage is likely an underestimate of the true size of the mouse pseudogene complement given the similarities between the human and mouse genomes. Thus, to get a more accurate estimate of the number of pseudogenes in the mouse genome, we used a combination of two automatic annotation pipelines: PseudoPipe [38] and RetroFinder [69]. PseudoPipe is the in-house comprehensive annotation pipeline that identifies and characterises pseudogenes based on their biotypes as either processed or duplicated. The automatic annotation workflow using PseudoPipe is summarized in **Figure 1B** and has been previously described in detail in [36, 37, 38]. Pseudopipe identified 22,811 mouse pseudogenes, of which 14,084 are present in autosomal chromosomes; this is comparable with previous reports in human (**Table S1**). RetroFinder is a computational annotation pipeline focused on identifying retrotransposed genes and pseudogenes. Using RetroFinder, we were able to annotate 18,467 and 15,474 processed pseudogenes in mouse and human, respectively. There was good overlap between the two automatic identification pipelines with respect to the number of processed pseudogenes present in both organisms (**Table S1**).

#### Mouse strain annotation

The mouse strain pseudogene annotation workflow is summarized in **Figure 1B**. The protein-coding input set contains the conserved protein-coding genes between each mouse strain and the reference genome. The number of shared transcripts follows an evolutionary trend with more distant strains having a smaller number of common protein-coding genes with the reference genome compared with more closely related laboratory strains. PseudoPipe was run with the strain conserved protein set as shown in **Figure 1B**. Next, we used the HAL tools package [70] to lift over the manually annotated pseudogenes from the mouse reference genome onto each strain using the UCSC multi-strain sequence alignments. We merged the two annotation sets using BEDTools [71] with a 1bp minimum overlap requirement. We extended the predicted boundaries to maximise the overlap and to ensure full annotation of the pseudogene transcript. Finally, we manually inspected the resultant annotation set in order to eliminate all potential false positives (e.g., pseudogene calls larger than 5Kb or smaller than 100bp, with poor protein-coding gene query similarity and coverage).

Next, we estimated the total number of pseudogenes in each strain.

Given the close evolutionary relationship between the mouse reference strain C57BL/6J and the laboratory “reference-like” strain C57BL/6NJ, we expect that given the same genome assembly quality and protein-coding annotation, the two strains will exhibit a similar number of pseudogenes and protein-coding genes. Further, the differences between C57BL/6NJ and the reference genome will give an indication of the quality of the laboratory strains’ annotation. Thus, we regarded C57BL/6NJ as a calibration strain.

To compute the number of pseudogenes in a particular strain based on the PseudoPipe annotation pipeline, we assumed that the correct number of pseudogenes is related to the number of input protein-coding transcripts. Moreover, observing that the number of conserved protein-coding transcripts between the mouse strains and the reference genome drops with increasing the evolutionary distance between the two, we worked on the premise that the actual total number of protein-coding gene transcripts should be constant across all mouse species. Thus, we defined a protein-coding transcript deflation factor as follows:

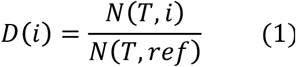

Where *N(T,i)* is the number of protein-coding transcripts in strain *i* that are used as input, and *N(T,ref)* is the number of protein-coding transcripts in the reference genome.

Next, in order to get a realistic estimate of the total number of pseudogenes in each strain based on Pseudopipe annotations, we needed to correct for strain quality by considering how the deflation affects the number of pseudogenes in the calibration strain, as this number should be the same as the reference. We computed the calibration strain correction factor as follows:

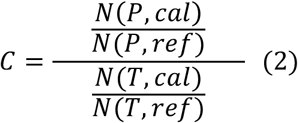

Where *N(P,cal)* is the number of PseudoPipe-annotated pseudogenes in the C57BL/6NJ calibration strain, *N(P,ref)* is the number of PseudoPipe pseudogenes annotated in the reference genome, *N(T,cal)* is the number of protein-coding transcripts used as input in the calibration strain, and *N(T,ref)* is the number of protein-coding transcripts used as input in the reference genome.

Thus, using the information from both the deflation factor and calibration strain correction factor we were able to estimate the number of pseudogenes in each strain based on the initial PseudoPipe pipeline output as follows:

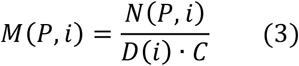

Where *N(P,i)* is the number of pseudogenes in strain *i*, *D(i)* is the deflation factor for strain *i*, and *C* is the calibration correction factor.

#### Unitary Pseudogene Annotation Pipeline

We modified PseudoPipe to allow cross-strain and cross-species protein-coding inputs. We annotated cross-organism pseudogenes as shown in **Figure 1B**. We define “Functional organism” as the genome providing the protein-coding information and thus containing a working copy of the element of interest; “Non-functional” organism denotes the genome queried for unitary pseudogene presence. The resulting data set was subjected to a number of filters such as removal of previously known pseudogenes, removal of pseudogenes with parents that have orthologs in the annotated species, removal of pseudogenes that overlap with annotated protein-coding and ncRNAs loci, and removal of pseudogenes shorter than 100bp. The filtered PseudoPipe set was intersected with the liftover of the protein-coding annotation from the functional organism using BEDTools [71] with a minimum of 1bp overlap required. The intersection set was further refined by flagging protein-coding genes that have functional relatives (paralogs) in the non-functional organism. The remaining matches were subjected to manual inspection of the alignment.

### Conservation and divergence in pseudogene complements

#### Pangenome data set generation

We performed an “all against all” liftover of pseudogene annotation using the HAL tools package and the UCSC multi-strain sequence alignment tool. Each liftover was intersected with the known strain annotation, and all of the entries that matched protein-coding genes or ncRNAs were removed. The resulting set was further filtered for conservation of pseudogene Ensembl ID (where available; Level 1 and 2 pseudogenes), parent gene identity, pseudogene locus (reciprocal overlap of 90% or higher), pseudogene biotype, pseudogene length, and pseudogene structure.

Next, we integrated all filtered binary mappings in a master pan-strain set. The common entries were collapsed into a unique pangenome pseudogene reference. We obtained 49,262 pangenome pseudogenes. A total of 1,158 pangenome entries were multi-matching across strains.

The number of strain-specific pseudogenes was calculated as the difference between the total number of pseudogenes and the number of pseudogenes with at least one identified ortholog in another strain/sub-species. The high specificity and accuracy in annotating orthologs translates into high sensitivity in identifying strain-specific pseudogenes. Thus, the current number of strain-specific pseudogenes is an upper bound of the expected dataset size. To estimate the lower bound of the number of strain-specific pseudogenes, we relaxed the cut-off level in the conservation of pseudogene locus and sequence overlap (see **Figure SF8**). The lower the threshold, the larger the number of called orthologs and, consequently, a smaller number of strain-specific pseudogenes. The minimum number of expected strain-specific pseudogenes in the current dataset was calculated under the hypothesis that a strain-specific pseudogene will have 0% sequence overlap with any annotated elements in any of the other strains. Thus, there are a minimum of 295 strain unique pseudogenes on average in any of the 18 mouse genomes.

#### Phylogenetic analysis

Sequences of the 1,460 pseudogenes were randomly selected out of the total 2,925 conserved pseudogenes in the 18 mouse strains; this accounts for approximately 50% of the total number of conserved pseudogenes. For each of the 18 mouse genomes, the extracted sequences were concatenated into a strain-specific contig (supergene). The order of the pseudogene sequences was kept the same in all 18 contigs. Preserving the same order of pseudogenes or protein-coding genes across all strains eliminates any potential bias. Also by selecting samples from the conserved pseudogenes set that are randomly distributed across the genome we are able to by-pass any potential bias introduced by the laboratory strain mosaicism. Thus, the resulting phylogeny depends only on sequence evolution. The 18 supergenes were subjected to multi-sequence alignment using MUSCLE aligner [72] under standard conditions. Similarly, the sequences of parent protein-coding genes of the 1,460 pseudogenes were assembled into a strain-specific sequence and aligned using MUSCLE. The tree was generated with PhyML using the Tamura-Nei genetic distance model and simultaneous Nearest Neighbor Interchange build method with *Mus Pahari* as the outgroup [73]. For both the pseudogene and parent gene, tree alignment and tree construction was done on the nucleotide level.

### Genome evolution and plasticity

#### Genome mappability maps

We created mappability maps for the mouse reference genome and the 18 mouse strains using the GEM library [74]. The workflow is composed of indexing the genome using gem-indexer, followed by creation of the map using a window of 75 nucleotides under the following conditions: -m 0.02 -T 2.

#### Parent gene expression analysis

RNA-seq mouse tissue data was obtained from ENCODE. The complete list of experiments used is available in **Table S8**. We estimated the expression levels of the pseudogene parent protein-coding genes using a workflow involving the following steps: filtering the protein-coding genes for uniquely mappable regions longer than 100bp, mapping reads using TopHat2 [75], selecting high-quality mapped reads with a quality score higher than 30 using samtools [76], and calculating the expression of FPKM levels using Cufflinks [77]. Transcriptional activity of pseudogene parent genes during early embryonic development was investigated using RNA-seq data as processed and described in [48]. Raw sequencing data and processed data containing FPKM levels at each embryonic stage are available on the SRA under Series GSE66582.

#### Transposable elements analysis

TEs in human and mouse reference genomes were informed from the RepeatMasker library Repbase 21.11 and using RepeatMasker 3.2.8 [78]. We extracted all four major groups of repeats, SINE, LINE, LTR, and DNA, and identified all the processed pseudogenes associated with L1 elements. Next, we binned the L1 annotated pseudogenes into age groups based on their sequence similarity to the parent gene, with younger elements exhibiting a higher sequence similarity and older elements showing a large sequence divergence when compared to the functional gene counterparts.

### Gene ontology and Pfam analysis

Linking of GO terms to the pseudogene parent genes was conducted using the R package biomaRt [79, 80]. Visualization of shared and distinct GO term sets among the strains was done using the R package UpSetR [81]. Enrichment of GO terms among the pseudogene parent genes and clustering of mouse strains based on similar enrichment profiles was performed using the goSTAG software package [82]. Semantic clustering of the GO terms was done with OntologyX packages [83]. Parent genes were labeled with both strain and biotype information in order to better evaluate differences in the pseudogene complements based on their mechanism of creation.

Analysis of the Pfam representation in the pseudogene complements was performed as previously described in [84] and focused on associating the pseudogene with the protein family of its parent gene.

### Gene essentiality enrichment analysis

Lists of essential and non-essential genes were compiled using data from the MGI database and recent work from the International Mouse Phenotyping Consortium [85]. The non-essential gene set with Ensembl identifiers contained 4,736 genes compared to 3,263 essential genes.

In order to evaluate the impact of parent gene status on the probability of a gene being essential while controlling for transcription, we fit a linear probability model and a probit model for the probability that a gene is essential given its transcription level and parent gene status using the StatsModels package in Python. The linear probability model fits an ordinary least squares regression of gene essentiality on parent gene status and transcription level. Although the linear probability model generally estimates relationships well close to the mean of the independent variables, it often loses explanatory power at low and high values of these variables. Because of this deficiency, we also used the probit model, which is similar to the linear probability model but instead fits the data to a cumulative Gaussian distribution. Around the mean values, we found that parent gene status increases the probability of essentiality by around 20% in both models.

### Pseudogene transcription

We estimated the pseudogene transcription levels for the mouse reference in 18 adult tissues following a similar protocol as the one described earlier for calculating the expression of protein-coding genes. we have successfully used this method in the past [37] using ENCODE RNA-seq data (**Table S8**). The pseudogene sequences were filtered for uniquely mappable exon regions longer than 100 bp. Next the RNA-seq raw data was mapped using TopHat and the mapped reads were filtered for quality scores higher than 30. The resulting alignments were quantified using Cufflinks. A pseudogene was considered transcribed if it had an FPKM larger than 3.3 in accord with previous studies [37].

RNA-seq data from mouse adult brain was obtained from the Mouse Genome Project for 12 laboratory and four wild-derived strains (ftp://ftp-mouse.sanger.ac.uk/REL-1509-Assembly-RNA-Seq, sanger experiment, last accessed on 21.08.2017). Next, we created mappability maps for each of the 16 mouse strains’ genomes and selected only the pseudogene exons in uniquely mappable regions that were longer than 100bp for further transcription analysis. The pseudogene transcription levels in mouse strains were estimated using a similar workflow as described above. The transcription cutoff level was set to 1.

### Mouse pseudogene resource

All of the annotation data produced in the analysis was collected and made available online through mouse.pseudogene.org. Pseudogene annotation information encompasses the genomic context of each pseudogene, its parent gene and transcript Ensembl IDs, the corresponding mouse reference pseudogene Ensembl ID, the level of confidence in the pseudogene as a function of agreement between manual and automated annotation pipelines, and the pseudogene biotype.

Information on the cross-strain comparison of pseudogenes is derived from the liftover of pseudogene annotations from one strain onto another and subsequent intersection with that strain’s native annotations. The database provides liftover annotations and information about intersections between the liftover and native annotations. Furthermore, homology information provides links between the well-characterised mouse strain collection.

Links between the annotated pseudogenes, their parent genes, and relevant functional and phenotypic information help inform biological relevance. In the database, the Ensembl ID associated with each parent gene is linked to the appropriate MGI gene symbol, which serves as a common identifier to connect to the phenotypic information. These datasets include information on gene essentiality, Pfam families, GO terms, and transcriptional activity.

## Acknowledgements

This project was supported by the Wellcome Trust (grant numbers WT108749/Z/15/Z, WT098051, WT202878/Z/16/Z and WT202878/B/16/Z), Cancer Research UK (20412), the European Research Council (615584), and the European Molecular Biology Laboratory. The research leading to these results has received funding from the European Union’s Seventh Framework Programme (FP7/2007- 2013) under grant agreement HEALTH-F4-2010-241504 (EURATRANS).

